# Activating soluble adenylyl cyclase protects mitochondria, rescues retinal ganglion cells, and ameliorates visual dysfunction caused by oxidative stress

**DOI:** 10.1101/2024.03.04.583371

**Authors:** Tonking Bastola, Guy A. Perkins, Viet Anh Nguyen Huu, Saeyeon Ju, Keun-Young Kim, Ziyao Shen, Dorota Skowronska-Krawczyk, Robert N. Weinreb, Won-Kyu Ju

## Abstract

Oxidative stress is a key factor causing mitochondrial dysfunction and retinal ganglion cell (RGC) death in glaucomatous neurodegeneration. The cyclic adenosine monophosphate (cAMP)/protein kinase A (PKA) signaling pathway is involved in mitochondrial protection, promoting RGC survival. Soluble adenylyl cyclase (sAC) is one of the key regulators of the cAMP/PKA signaling pathway. However, the precise molecular mechanisms underlying the sAC-mediated signaling pathway and mitochondrial protection in RGCs that counter oxidative stress are not well characterized. Here, we demonstrate that sAC plays a critical role in protecting RGC mitochondria from oxidative stress. Using mouse models of oxidative stress, we found that activating sAC protected RGCs, blocked AMP-activated protein kinase activation, inhibited glial activation, and improved visual function. Moreover, we found that this is the result of preserving mitochondrial dynamics (fusion and fission), promoting mitochondrial bioenergetics and biogenesis, and preventing metabolic stress and apoptotic cell death in a paraquat oxidative stress model. Notably, sAC activation ameliorated mitochondrial dysfunction in RGCs by enhancing mitochondrial biogenesis, preserving mitochondrial structure, and increasing ATP production in oxidatively stressed RGCs. These findings suggest that activating sAC enhances the mitochondrial structure and function in RGCs to counter oxidative stress, consequently promoting RGC protection. We propose that modulation of the sAC-mediated signaling pathway has therapeutic potential acting on RGC mitochondria for treating glaucoma and other retinal diseases.

## INTRODUCTION

Glaucoma is a chronic multifactorial disease with multiple risk factors, including intraocular pressure (IOP) and oxidative stress, and leads to irreversible blindness on a global scale^1^. Findings from our research group and other studies strongly suggest that compromised mitochondrial function and metabolic stress by glaucomatous insulting factors, such as elevated IOP and oxidative stress, play a crucial role in the degeneration of retinal ganglion cell (RGC) somas and axons in experimental glaucoma^2, 3^; this indicates that impaired mitochondrial network and function are strongly linked to glaucoma pathogenesis^2, 3^. Although the significance of mitochondrial dysfunction and loss in the context of disease is widely acknowledged, the understanding of molecular mechanisms underlying oxidative stress, mitochondrial health, or the prevention of mitochondrial dysfunction in glaucoma remains elusive.

Cyclic adenosine monophosphate (cAMP) serves as a widely distributed second messenger in the central nervous system (CNS) to facilitate signal transduction^4^. Upon activation, the synthesis and degradation of cAMP are intricately controlled by adenylyl cyclases (ACs) and cyclic nucleotide phosphodiesterases^5^. The activity of cAMP is modulated by diverse effectors, including cAMP-dependent protein kinase A (PKA)^6^, guanine-nucleotide exchange proteins^7^, and cyclic-nucleotide-gated ion channels^8^. Within the cAMP signaling pathway, ACs are crucial regulators, functioning as enzymes responsible for synthesizing cAMP from adenosine 5′-triphosphate (ATP). At present, there exist ten individual AC genes (AC1-10), encompassing nine mammalian transmembrane ACs (AC1-9) and a soluble AC (sAC; AC10)^9^.

Current evidence indicates that the sAC/cAMP/PKA signaling pathway is linked to mitochondrial protection, ultimately promoting cell survival in neurons, notably in RGCs^10, 11^. The sAC modulates the cAMP activity within the mitochondrial matrix^12, 13^, and two mitochondrial cAMP pools influenced by sAC are associated with the outer mitochondrial membrane and the intermembrane space of mitochondria^14, 15^. Considering that the activation of the sAC-mediated cAMP/PKA signaling pathway is crucial for boosting the survival of RGCs and promoting axon growth^11, 16^, RGC protection from elevated IOP and oxidative stress could serve as a therapeutic approach in addressing glaucomatous neurodegeneration. Importantly, oxidative stress is a critical causative factor in mitochondrial dysfunction and RGC death in glaucoma.

In this study, we highlight the critical role of sAC in RGC mitochondria protection from oxidative stress. We demonstrate that activating sAC protects RGCs and ameliorates the degrading of visual function due to oxidative stress by preserving mitochondrial dynamics (fusion and fission), promoting mitochondrial bioenergetics and biogenesis, and preventing metabolic stress and apoptotic cell death in experimental mouse models of oxidative stress. Our findings indicate the therapeutic potential of sAC in preventing impaired mitochondrial dynamics and function, particularly in the context of oxidative stress-mediated glaucomatous neurodegeneration.

## RESULTS

### sAC activation protects RGCs by increasing mitochondrial biogenesis and inhibiting Bax activation in the retina to counter oxidative stress

To determine whether sAC activation protects RGCs from retinal oxidative stress, mice were administered regular drinking water or drinking water containing 150 mM sodium bicarbonate (NaHCO_3_) at 1 week before a transient ischemia-reperfusion incidence by acute IOP elevation (Fig. 1A). We found that elevated IOP-induced oxidative stress significantly induced RGC loss in the retina compared with non-ischemic control retina (Fig. 1B and C). In contrast, we remarkably observed that sAC activation by NaHCO_3_ administration significantly promoted RGC survival in ischemic retinas (Fig. 1B and C). sAC activation enhances peroxisome proliferator-activated receptor-gamma coactivator (PGC1-α), a key regulator of energy metabolism and potential stimulator of mitochondrial biogenesis^17, 18^. We found that elevated IOP-induced oxidative stress significantly reduced PGC1-α protein expression in the retina. In contrast, sAC activation remarkably restored the expression level of PGC1-α protein in the ischemic retina (Fig. 1D). Moreover, we observed that sAC activation significantly reduced the expression level of active BAX protein in the ischemic retina (Fig. 1D). In addition, NaHCO_3_ administration significantly increased cAMP immunoreactivity in the retina (Fig. S1).

**Figure 1.**
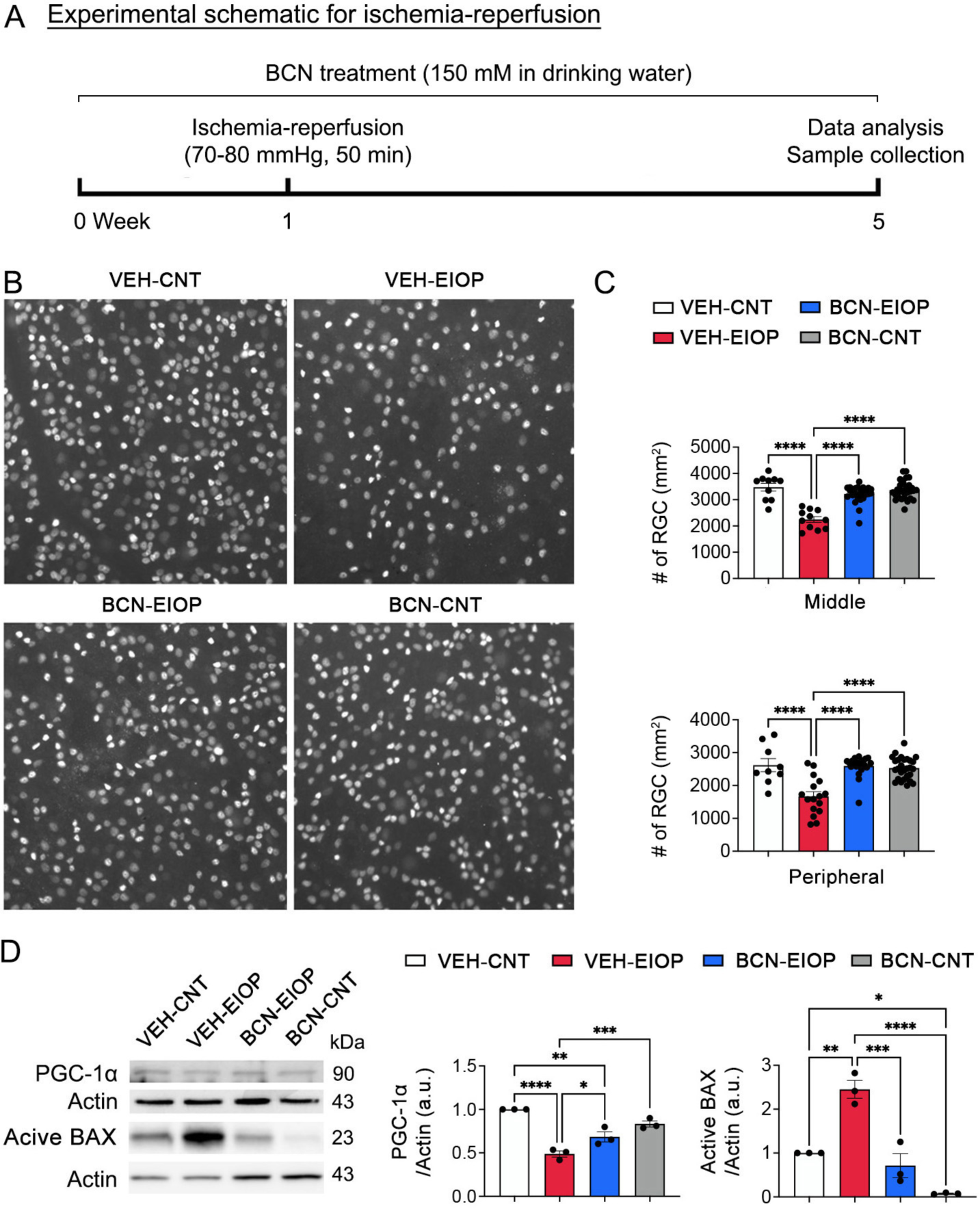
sAC activation protects RGCs by increasing mitochondrial biogenesis and inhibiting Bax activation in the oxidatively stressed retina. (*A*) Diagram for control and bicarbonate (BCN, 150 mM) administration, sample collection and data analysis. Four-month-old C57BL/6J mice were administered either regular drinking water or drinking water containing BCN for 5 weeks. The drinking water or BCN in drinking water were given weekly. (*B*) Retinal whole-mount immunohistochemistry using BRN3a antibody. Representative images showed BRN3a-positive RGCs in the middle and peripheral areas of the retinas from each group. (*C*) Quantitative analysis of RGC survival. *N* = 5 mice per group. (*D*) PGC1-α and active BAX protein expression in the retina. *N* = 3 mice per group. Error bars represent SEM. Statistical significance determined using one-way ANOVA test. \**P* < 0.05; \*\**P* < 0.01; \*\*\**P* < 0.001; \*\*\*\**P* < 0.0001. Scale bar: 50 μm. BCN, bicarbonate; CNT, control; EIOP, elevated intraocular pressure. See also Figure S1.

### sAC activation restores visual function that had been reduced by oxidative stress

To test the effect of sAC activation on visual function recovery from oxidative stress, mice were administered regular drinking water or drinking water containing 150 mM NaHCO_3_ at 1 week before treatment of paraquat (PQ, 15 mg/kg), which is an oxidative stress inducer^19^, increasing mitochondrial oxidative damage (Fig. 2A). In conditions of oxidative stress, we observed a noteworthy decline in visual acuity, indicated by reduced spatial frequency and visual evoked potential (VEP) P1-N1 potentials, along with increased latency in mice, as measured through optomotor response and VEP in mice (Fig. 2B and C). In contrast, sAC activation remarkably restored spatial frequency and VEP P1-N1 potentials in mice exposed to oxidative stress (Fig. 2B and C). However, there were no statistically significant differences in latency in mice exposed to oxidative stress (Fig. 2C).

**Figure 2.**
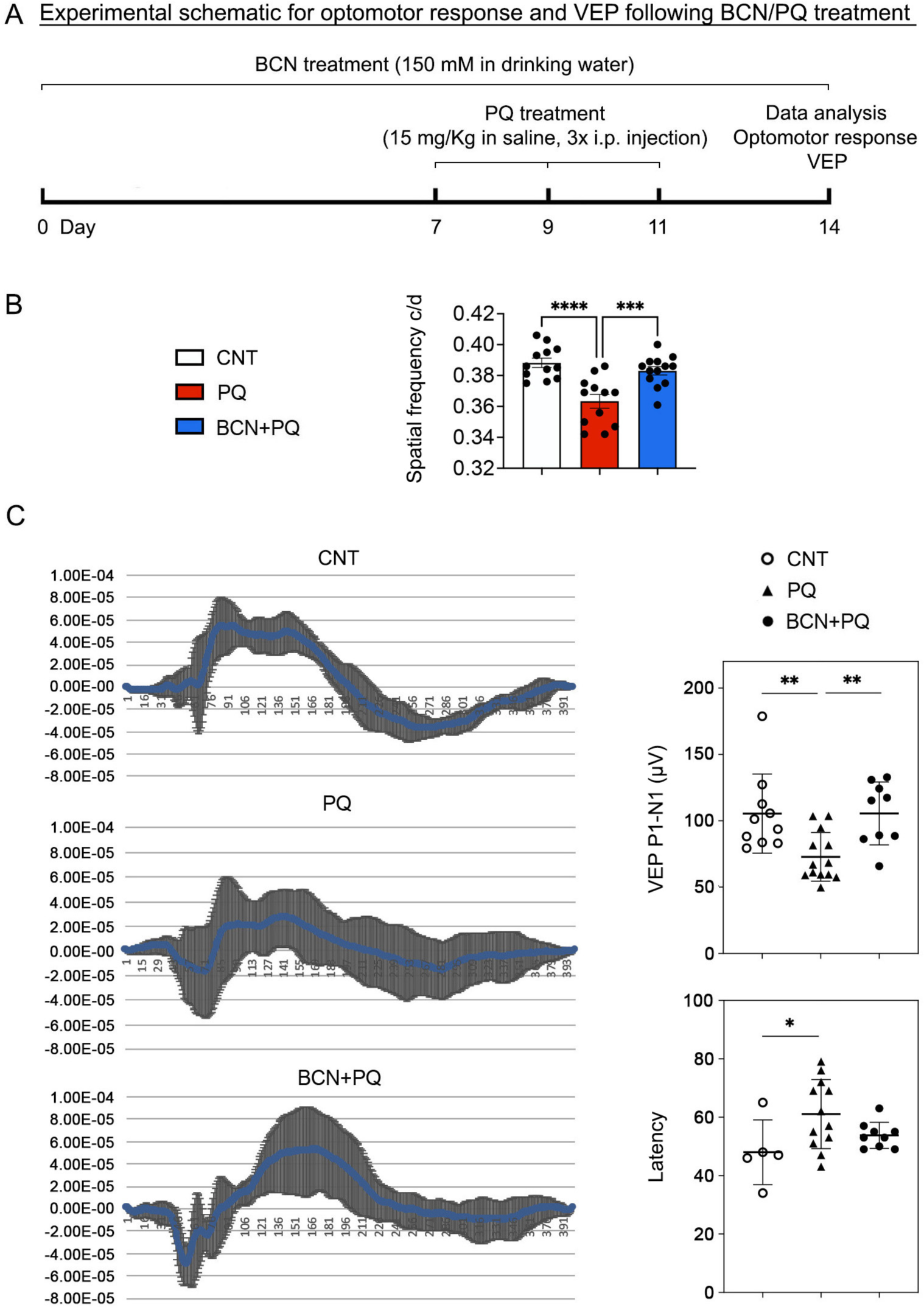
sAC activation restores visual function reduced by oxidative stress. (*A*) Diagram for control and bicarbonate (BCN, 150 mM) administration, sample collection and data analysis. Four-month-old C57BL/6J mice were administered either regular drinking water or drinking water containing BCN for 2 weeks. The drinking water or BCN in drinking water were given weekly. The mice received PQ (15 mg/Kg) by i.p. injection three times for 1 week after 1 week of BCN pretreatment. (*B*) Visual function test in mice by optomotor response analyses. *N* = 8 to 9 mice. (*C*) Total recordings of VEP responses among groups. (*D*) Visual function test in mice by VEP analyses. *N* = 9 to 10 mice. Error bars represent SEM. Statistical significance determined using one-way ANOVA test. \**P* < 0.05; \*\**P* < 0.01; \*\*\**P* < 0.001; \*\*\*\**P* < 0.0001. BCN, bicarbonate; CNT, control; EIOP, elevated intraocular pressure; PQ, paraquat; VEP, visual evoked potential.

### sAC activation inhibits glial activation, p38 phosphorylation and BAX activation in the retina to counter oxidative stress

Since oxidative stress induces glial activation (astrocytes and microglial cells) in the retina^20^, we determined whether sAC activation inhibits glial activation in the retina to oppose oxidative stress (Fig. 3A). We observed a significant increase of GFAP and IBA-1 expression in the retina exposed to oxidative stress (Fig. 3B. However, sAC activation significantly decreased the expression levels of GFAP and IBA-1 in the retina to oppose oxidative stress (Fig. 3B). Oxidative stress is associated with the activation of mitogen-activated protein kinases (MAPKs), including p38, and cell death in the retina^21^. We remarkably found a significant increase of phospho-p38 (pp38) expression in the retina exposed to oxidative stress (Fig. 3C). In contrast, sAC activation significantly decreased the expression level of pp38 in the retinas exposed to oxidative stress (Fig. 3C). Oxidative stress triggers BAX activation and induces apoptotic cell death in the retina^22, 23, 24^. We observed a significant expression of active BAX but a decreased pattern of BCL-xL) in the retina exposed to oxidative stress (Fig. 3D). However, sAC activation significantly decreased the expression level of active BAX and showed an increased pattern of BCL-xL expression in the retina exposed to oxidative stress (Fig. 3D).

**Figure 3.**
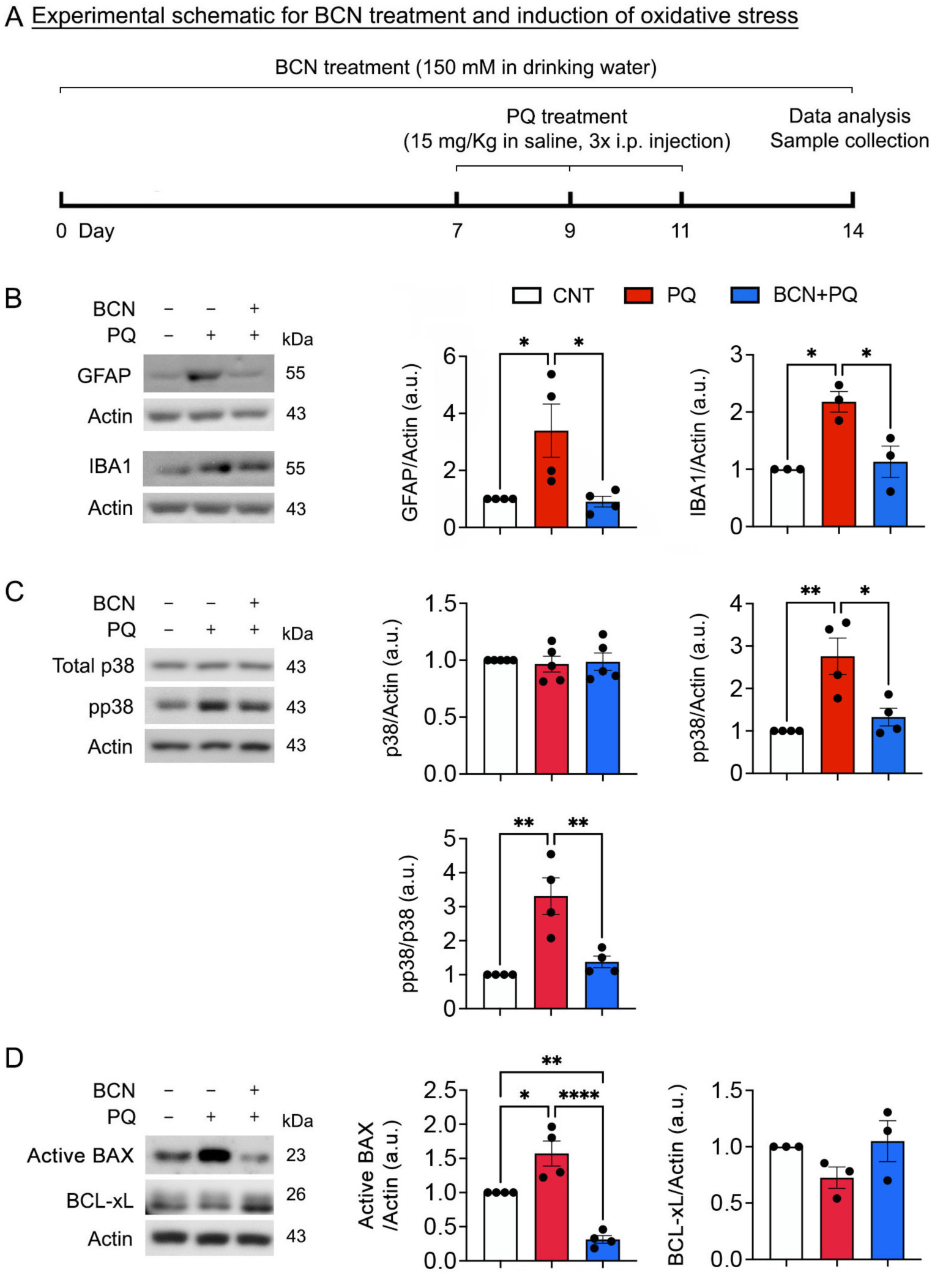
sAC activation inhibits glial activation, and MAPK and BAX activation in the oxidatively stressed retina. (*A*) Diagram for control and bicarbonate (BCN, 150 mM) administration, sample collection and data analysis. (*B*) GFAP and IBA1 protein expression in the retina. *N* = 3 mice per group. (*C*) Total p38 and phospho-p38 (pp38) protein expression in the retina. *N* = 3 mice per group. (*D*) Active BAX and BCL-xL protein expression in the retina. *N* = 3 to 4 mice per group. Error bars represent SEM. Statistical significance determined using one-way ANOVA test. \**P* < 0.05; \*\**P* < 0.01; \*\*\*\**P* < 0.0001.

### sAC activation enhances the expression of AKAP1 and PKAα, phosphorylation of DRP1 at Ser637 and GSK3β at Ser9, as well as mitochondrial biogenesis and OXPHOS in the retina subjected to oxidative stress

The sAC-cAMP-PKA signaling axis contributes to the phosphorylation of mitochondrial proteins, modulation of the oxidative phosphorylation (OXPHOS) system, and regulation of metabolic enzymes^25, 26^. We determined whether sAC activation by NaHCO_3_ treatment modulates phosphorylation of mitochondria-related proteins, which are associated with mitochondrial dynamics, in the retina exposed to oxidative stress (Fig. 4A). We found that oxidative stress significantly increased total dynamin-related protein 1 (DRP1) protein expression but triggered dephosphorylation of DRP1 at serine 637 (S637) in the retina at 1 week (Fig. 4B). However, phosphorylation of DRP1 at serine 616 (S616) was not changed in the retina exposed to oxidative stress (Fig. 4B). In parallel, we observed that oxidative stress significantly decreased A-kainase anchoring protein 1 (AKAP1) and PKAα protein expression in the retina (Fig. 4B). In contrast, we found that sAC activation significantly restored AKAP1 and PKAα protein expression and phosphorylation of DRP1 S637 and DRP1 S616 in the retina under oxidative stress (Fig. 4B and C, Fig. S2A). Interestingly, we also found that sAC activation did not significantly change the expression levels of the mitochondrial fusion proteins, optic atrophy type 1 (OPA1) and mitofusin (MFN1)/2 (Fig. S3). Because DRP1 is also regulated by glycogen synthase kinase 3 β (GSK3β)^27, 28, 29^, we further determined whether sAC activation modulates phosphorylation of GSK3β at Serine 9 (S9) in the retina exposed to oxidative stress. We found that oxidative stress significantly induced dephosphorylation of GSK3β S9 in the retina (Fig. 4D, Fig. S2B). Based on our previous findings of decreased DRP1 S637 phosphorylation in the retina of AKAP1 knockout (*AKAP1^−/−^*) mice^30^, we also found that AKAP1 deficiency significantly induced dephosphorylation of GSK3β S9 in the retina of *AKAP1^−/−^* mice (Fig. S4). Impairment of OXPHOS dysfunction is linked to patients with primary open-angle glaucoma^31^ and there is supporting evidence indicating that OXPHOS serves as the primary location for the production of mitochondrial superoxide in the presence of PQ^19^. Because the sAC/cAMP pathway is associated with mitochondrial biogenesis and OXPHOS function^10, 12, 32, 33, 34^, we next examined whether sAC activation promotes expression levels of PGC-1α and mitochondrial transcription factor A (TFAM) proteins, and OXPHOS complexes in the retina affected by oxidative stress. We found that oxidative stress significantly decreased PGC-1α and TFAM protein expression in the retina (Fig. 4E). In contrast, sAC activation increased PGC-1α and TFAM protein expression in the stressed retina (Fig. 4E). In addition, we observed that sAC activation enhanced OXPHOS complexes I and IV in the retina against oxidative stress (Fig. 4F).

**Figure 4.**
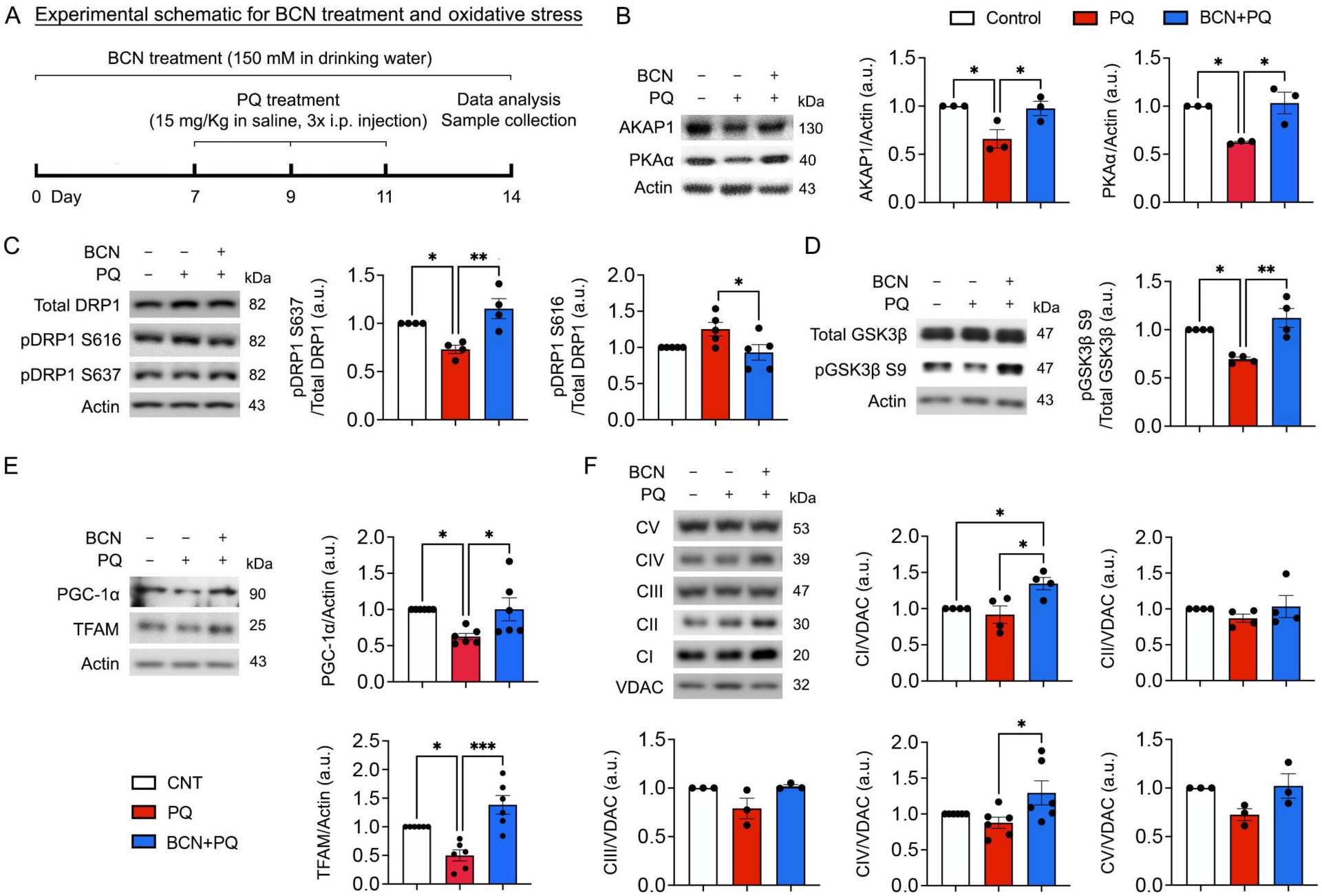
sAC activation enhances phosphorylation of DRP1 and GSK3β, as well as induces mitochondrial biogenesis and increases OXPHOS complexes in the oxidatively stressed retina. (*A*) Diagram for control and bicarbonate (BCN, 150 mM) administration, sample collection and data analysis. (*B*) AKAP1 and PKAα protein expression in the retina. *N* = 3 mice per group. (*C*) Phospho-DRP1 S616 and phospho-DRP1 S637 protein expression in the retina. *N* = 3 mice per group. (*D*) Phospho-GSK3β S9 protein expression in the retina. *N* = 3 mice per group. (*E*) PGC1-α and TFAM protein expression in the retina. *N* = 3 mice per group. (*F*) OXPHOS complex protein expression in the retina. *N* = 3 to 5 mice per group. Error bars represent SEM. Statistical significance determined using one-way ANOVA test. \**P* < 0.05; \*\**P* < 0.01; \*\*\**P* < 0.001.

### sAC activation reduces mitochondrial stress by blocking AMPK activation in the retina subjected to oxidative stress

5’ AMP-activated protein kinase (AMPK) is a highly conserved energy sensor and its activation is associated with glaucomatous RGCs^35^. Inactivation of AMPK promotes RGC survival in experimental glaucoma^36^. We determined whether sAC activation inhibits AMPK activation in the retina under oxidative stress (Fig. 5A). We found a significant increase of pAMPK (Thr172) in the stressed retina (Fig. 5A B). In contrast, we observed that sAC activation significantly decreased pAMPK in the stressed retina (Fig. 5B). Consistent with an increased pAMPK level, pAMPK immunoreactivity was significantly increased in RGCs as well as in the inner retina layers and outer plexiform layer in the stressed retina (Fig. 5C and D). However, we remarkably observed that sAC activation significantly decreased pAMPK immunoreactivity in the stressed retina (Fig. 5C and D). Downregulation of sAC is linked to impaired mitochondrial clearance^37^. However, it is unknown whether sAC contributes to the regulation of mitophagy in the retina exposed to oxidative stress. We further determined whether sAC activation reduces mitophagy in the stressed retina. Oxidative stress significantly enhanced microtubule-associated protein 1A/1B-light chain 3-(LC3)-II protein expression but decreased p62 protein expression in the retina. In contrast, sAC activation significantly reduced LC3-II protein expression but increased p62 protein expression in the stressed retina (Fig. 5E).

**Figure 5.**
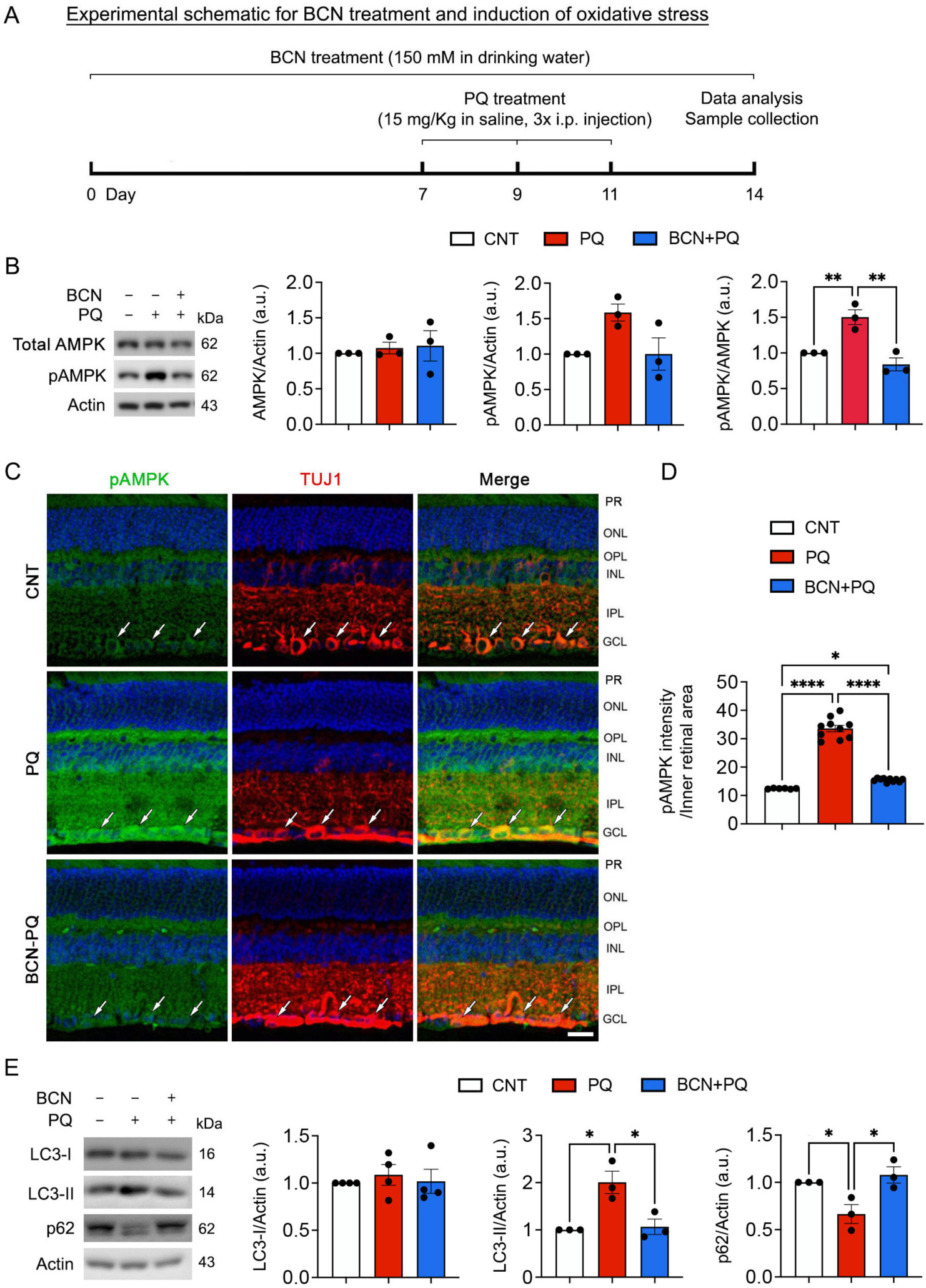
sAC activation promotes mitochondrial function and ameliorates AMPK activation in the oxidatively stressed retina. (*A*) Diagram for control and bicarbonate (BCN, 150 mM) administration, sample collection and data analysis. (*B*) Total AMPK and phospho-AMPK protein expression in the retina. *N* = 3 mice per group. (*C*) Representative images showed phospho-AMPK (green) and TUJ1 (red) immunoreactivities. (*D*) Phospho-AMPK immunoreactive intensity in the inner retina. *N* = 3 t mice per group. (*E*) LC3 and p62 protein expression in the retina. *N* = 3 to 4 mice per group. Error bars represent SEM. Statistical significance determined using one-way ANOVA test. \**P* < 0.05; \*\**P* < 0.01. Blue is Hoechst 33342 staining. Scale bar: 20 μm. GCL, ganglion cell layer; INL, inner nuclear layer; IPL, inner plexiform layer; ONL, outer nuclear layer; OPL, outer plexiform layer; PR, photoreceptor.

### sAC activation promotes mitochondrial biogenesis and ATP production in RGCs affected by oxidative stress

sAC/cAMP signaling contributes to neuronal cell survival and neurite outgrowth^11, 16^. To determine whether sAC activation protects RGCs and restores mitochondrial function that had been reduced by oxidative stress, we performed MitoTracker Red staining, a marker for mitochondria, and immunocytochemistry using an antibody for β-III-Tubulin (TUJ1), a marker for the RGC soma, axon and dendrites, at 1 day after PQ treatment (Fig. 6A). We remarkably found that MitoTracker Red signals and TUJ1-positive immunoreactivity were diminished in RGCs exposed to oxidative stress (Fig. 6B), suggesting that there was loss of neurites and mitochondria. However, sAC activation restored MitoTracker Red signals and TUJ1 immunoreactivity in the stressed RGCs (Fig. 6B). More importantly, we observed that sAC activation promoted neurite outgrowth in RGCs affected by oxidative stress (Fig. 6C).

**Figure 6.**
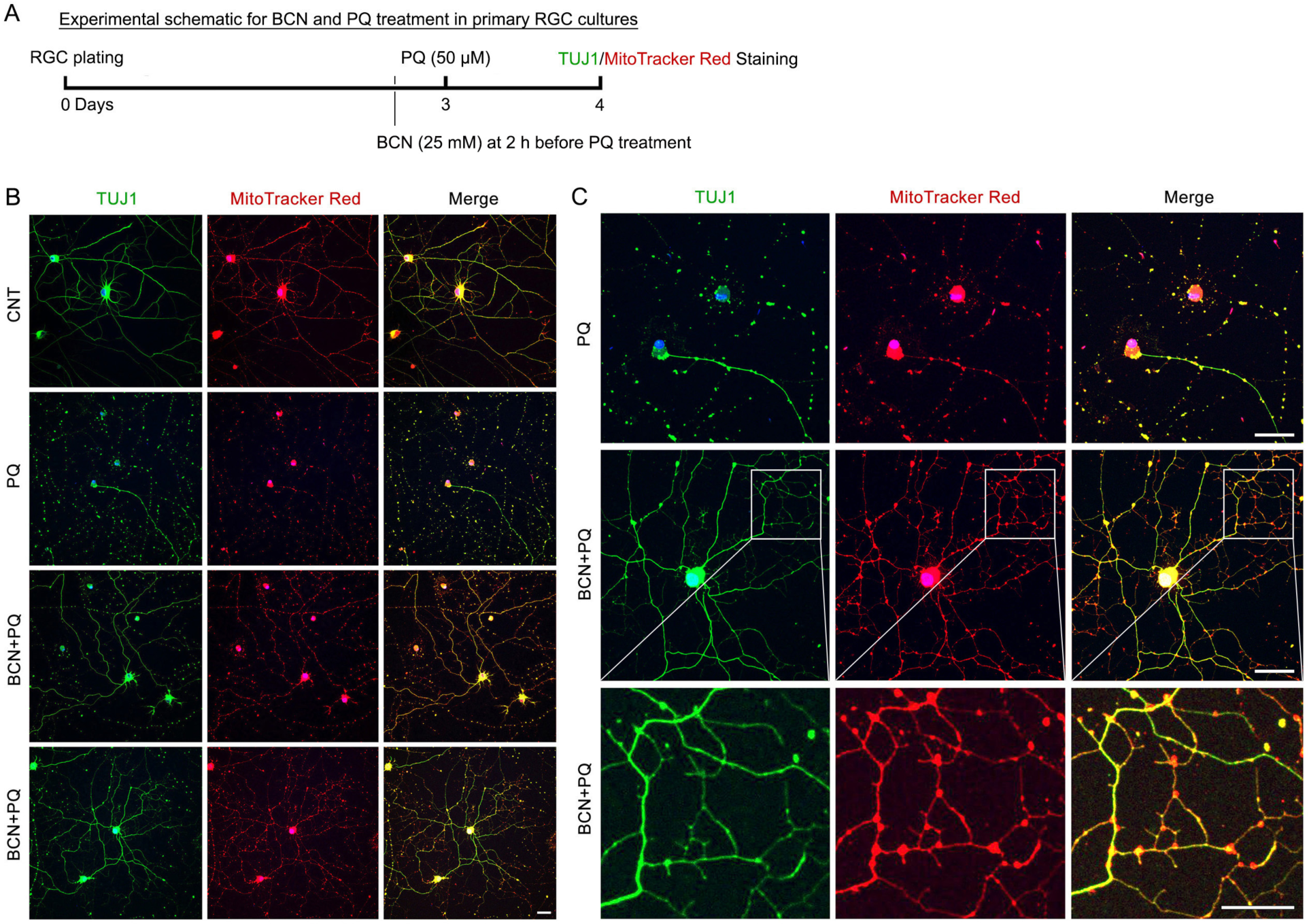
sAC activation promotes neurite outgrowth in oxidatively stressed RGCs. (*A*) Diagram for BCN (25 mM) and PQ (50 μM) treatment, sample collection and data analysis. Cultured RGCs were pretreated with BCN for 2 h before PQ treatment. PQ treatment lasted 24 h. (Band C) Representative images showed TUJ1 (green) immunoreactivity and MitoTracker staining (red) in cultured RGCs. Scale bar: 10 μm.

EM tomography showed that mitochondria were typically elongated with slightly condensed matrix (expanded cristae) in control RGCs (Figs. 7A and Ba-Bd, Movie 1). Control mitochondria were observed to be in a slightly condensed state likely indicating an increased rate of respiration (Fig. 7Ba). The tubular nature of control mitochondria was demonstrated by the 3D surface rendering of the outer mitochondrial membrane (OMM) (Fig. 7Bb). The surface-rendered volume also revealed the distribution and shape of the cristae (Fig. 7Bc and Bd). The cristae adopted a mostly lamellar shape (Fig. 7Bd). However, oxidative stress produced longer mitochondria, yet fewer in number, and abnormal mitochondrial membranes (Fig. 7Be-Bi, Movie 2). We observed an invaginated of the OMM to form an abnormal chamber (Fig. 7Be). The size and shape of the invagination was demonstrated by 3D surface-rendering (Fig. 7Bf). In addition, the surface-rendered volume showed the distribution and shape of the cristae (Fig. 7Bg), being lamellar in shape with some extending “fingers” towards the periphery. An example of a structural defect was seen in an invaginated portion of the OMM becoming vesiculated (Fig. 7Bh). In addition, one of the cristae close to the vesiculated OMM formed an abnormal tube-within-a-tube structure (Fig. 7Bi). Another structural defect was seen in the form of the OMM creating a completely vesiculated chamber within the mitochondrion and the tube-within-a-tube structure becoming separated from its parent crista, yet remained close to it as it extended through the volume (Fig. 7Bj,k,l). Remarkably, we found that sAC activation increased the crista density (Fig. 7Bm-Bp, Movie 3). In addition, the crista shape was roughly equally divided between lamellar (49 cristae) and tubular (60 cristae), yet the tubular cristae were arrayed around the mitochondrial periphery and altogether contained only 14% of the total cristae membrane surface area (Fig. 7Bq,r). In contrast, the lamellar cristae filled the center of the mitochondrion and were much larger (Fig. 7Bs,t).

**Figure 7.**
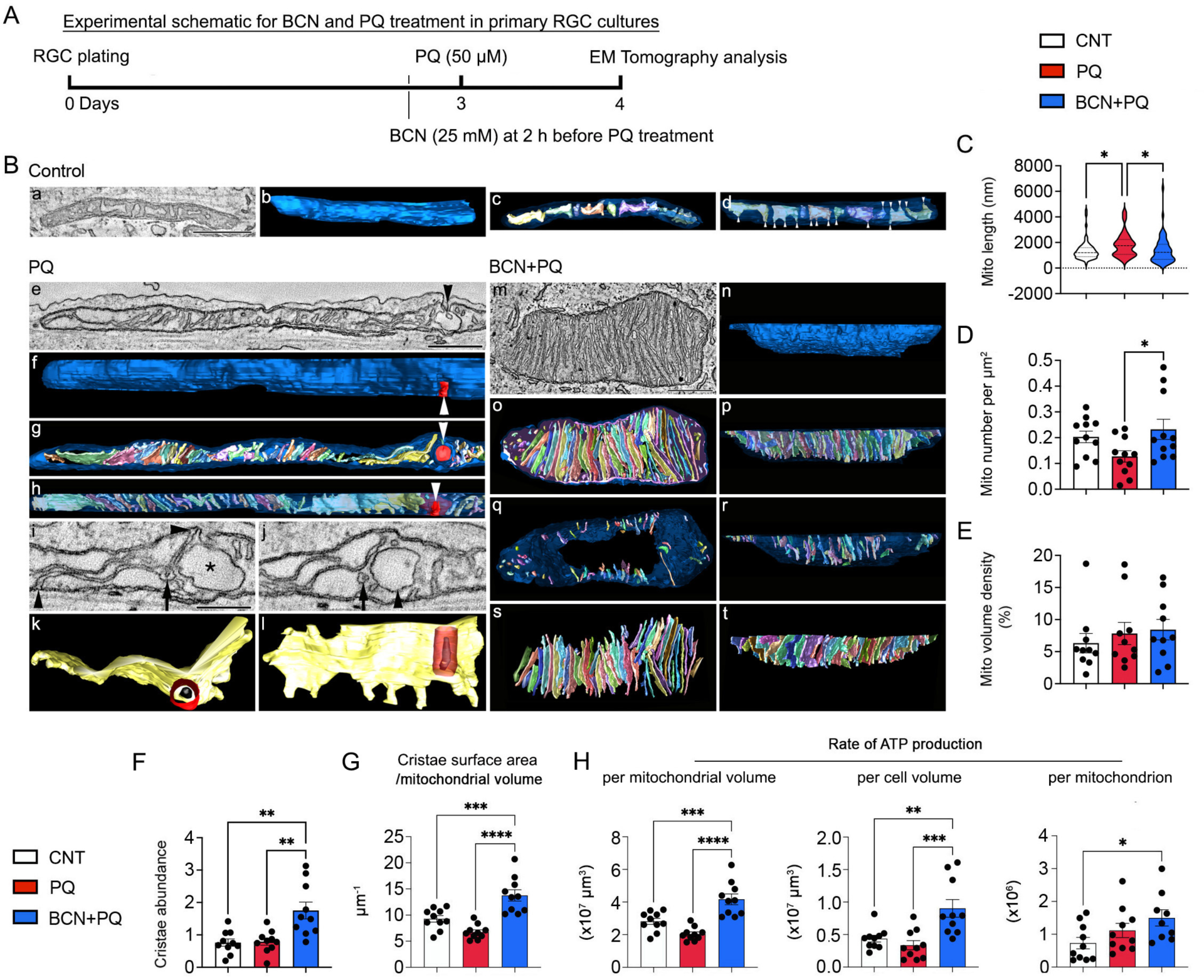
sAC activation promotes mitochondrial bioenergetic function in RGCs to counter oxidative stress. (*A*) Diagram for BCN (25 mM) and PQ (50 μM) treatment, sample collection and data analysis. BCN was added to cultured RGCs 2 h before PQ treatment. PQ treatment lasted 24 h. (*B*) **Control (a-d)**: Control mitochondria were typically elongated with slightly condensed matrix (expanded cristae) likely indicating an increased rate of respiration. **a** A 1.6-nm thick slice through the middle of a tomographic volume of a representative control mitochondrion. Scale = 500 nm. **b** Side view of the 3D surface-rendered mitochondrial outer membrane shown in blue to demonstrate the tubular nature of control mitochondria. **c** Top view of the surface-rendered volume showing the distribution and shape of the cristae (in an assortment of colors) with the mitochondrial outer membrane made translucent to better visualize the cristae. Nine cristae were present in this mitochondrion. **d** Side view of the surface-rendered volume showing the mostly lamellar shape of the cristae, but with many extending “fingers” (arrowheads) towards the periphery. **PQ (e-l)**: PQ treatment produced longer mitochondria, yet fewer in number and abnormal mitochondrial membranes. **e** A 2.0-nm thick slice through the middle of a tomographic volume of a PQ-treated mitochondrion. At one place, the mitochondrial outer membrane invaginated inwards to form a chamber (arrowhead). Scale = 500 nm. **f** Side view of the surface-rendered mitochondrial outer membrane shown in blue to demonstrate the size and shape of the invagination (arrowhead). **g** Top view of the volume showing the distribution and shape of the cristae. This mitochondrion had 35 cristae. Some of the cristae are seen to twist through the volume. The invaginated outer membrane is indicated by the arrowhead. **h** Side view of the surface-rendered volume showing the mostly lamellar shape of the cristae with some extending “fingers” towards the periphery. The invaginated outer membrane becomes vesiculated (arrowhead). **i** One of the cristae close to the vesiculated outer membrane (*) forms an abnormal tube-within-a-tube structure (arrow). Normal crista junctions are seen at both ends of this crista (arrowheads). Scale = 250 nm. **j** The outer membrane forms a completely vesiculated chamber within the mitochondrion (arrowhead) and the tube-within-a-tube structure (arrow) becomes separated from its parent crista, yet remained close to it as it extended through the volume. **k** Top view of the twisted crista emphasized in panels e and f showing the spatial relation of the tube-within-a-tube structure that extended from the crista (outer membrane: red, inner membrane: charcoal). **l** Side view of the same crista showing the extent of the outer membrane (red) and much smaller inner membrane (charcoal) of the abnormal tube-within-a-tube structure. The crista “fingers” extending towards the mitochondrial periphery are clearly seen. **BCN + PQ (m-r):** Treatment of PQ-exposed cells with BCN restored the number of mitochondria and their length to control levels, yet increased the crista density. **m** A 1.6-nm thick slice through the middle of a tomographic volume of a PQ+BCN mitochondrion that shows the dense packing of cristae. Scale = 500 nm. **n** Side view of the surface-rendered mitochondrial outer membrane. **o** Top view of the volume showing the distribution and shape of all 109 cristae. The 3D surface rendering further emphasized the high density of cristae. **p** Side view of the volume. The cristae were about equally divided between lamellar shape and tubular shape. **q** Top view of only the 60 tubular cristae, some of which were small. Interestingly, the tubular cristae were arrayed around the mitochondrial periphery and altogether contained only 14% of the total cristae membrane surface area. **r** Side view of the tubular cristae. **s** Top view of only the 49 lamellar cristae. These are much larger than the tubular cristae and occupy the central portion of the mitochondrial volume. **t** Side view of the lamellar cristae. (*C-H*) Measurements of structural and functional features of mitochondria. Statistical significance determined using a one-way ANOVA test. \**P* < 0.05; \*\**P* < 0.01; \*\*\**P* < 0.001; \*\*\*\**P* < 0.0001.

RGCs exposed to oxidative stress experienced significant structural changes. Mitochondrial length was increased (Fig. 7C, *n* = 60 mitochondria in 10 RGC somas). Mitochondrial number per cell area was decreased (Fig. 7D, *n* = 11 RGC somas). However, oxidative stress did not change mitochondrial volume density, cristae abundance, or crista density (defined as total cristae membrane surface area divided by the mitochondrial volume) (Fig. 7E-G, *n* = 10 RGC somas). Interestingly, we found that sAC activation significantly reduced mitochondrial length to a level similar to control mitochondria (Fig. 7C, *n* = 60 mitochondria in 10 RGC somas) and increased mitochondrial number, again to a level similar to control mitochondria in RGCs subjected to oxidative stress (Fig. 7D, *n* = 11 RGC somas). sAC increased cristae abundance and crista density even above control levels (Fig. 7F and G, *n* = 10 RGC somas). However, sAC activation did not change mitochondrial volume density (Fig. 7E, *n* = 10 RGC somas). We next performed energy calculations using a biophysical model based on 3D structure of OMM, a crista, and a crista junction ^38^. We observed no significant changes of the modeled rate of ATP production per mitochondrial volume, cell volume, or mitochondrion in stressed RGCs (Fig. 7H, *n* = 10 RGC somas). However, we remarkably found that sAC activation significantly increased the rate of ATP production per mitochondrial volume, cell volume and mitochondrion in RGCs (Fig. 7H, *n* = 10 RGC somas) subjected to oxidative stress.

### sAC activation promotes mitochondrial respiration in RGCs subjected to oxidative stress

Based on our findings of activating sAC-mediated enhancement of mitochondrial biogenesis and energy production, we examined the effect of sAC activation on mitochondrial activity and respiratory function in RGCs in response to oxidative stress. We measured mitochondrial activity and oxygen consumption rate (OCR) at 6 h after PQ treatment (Fig. 8A). We found that sAC activation significantly restored mitochondrial activity in RGCs under oxidative stress (Fig. 8B). Moreover, we remarkably found that sAC activation significantly increased basal and maximal respiration, and ATP-linked respiration in RGCs under oxidative stress (Fig. 8C and D).

**Figure 8.**
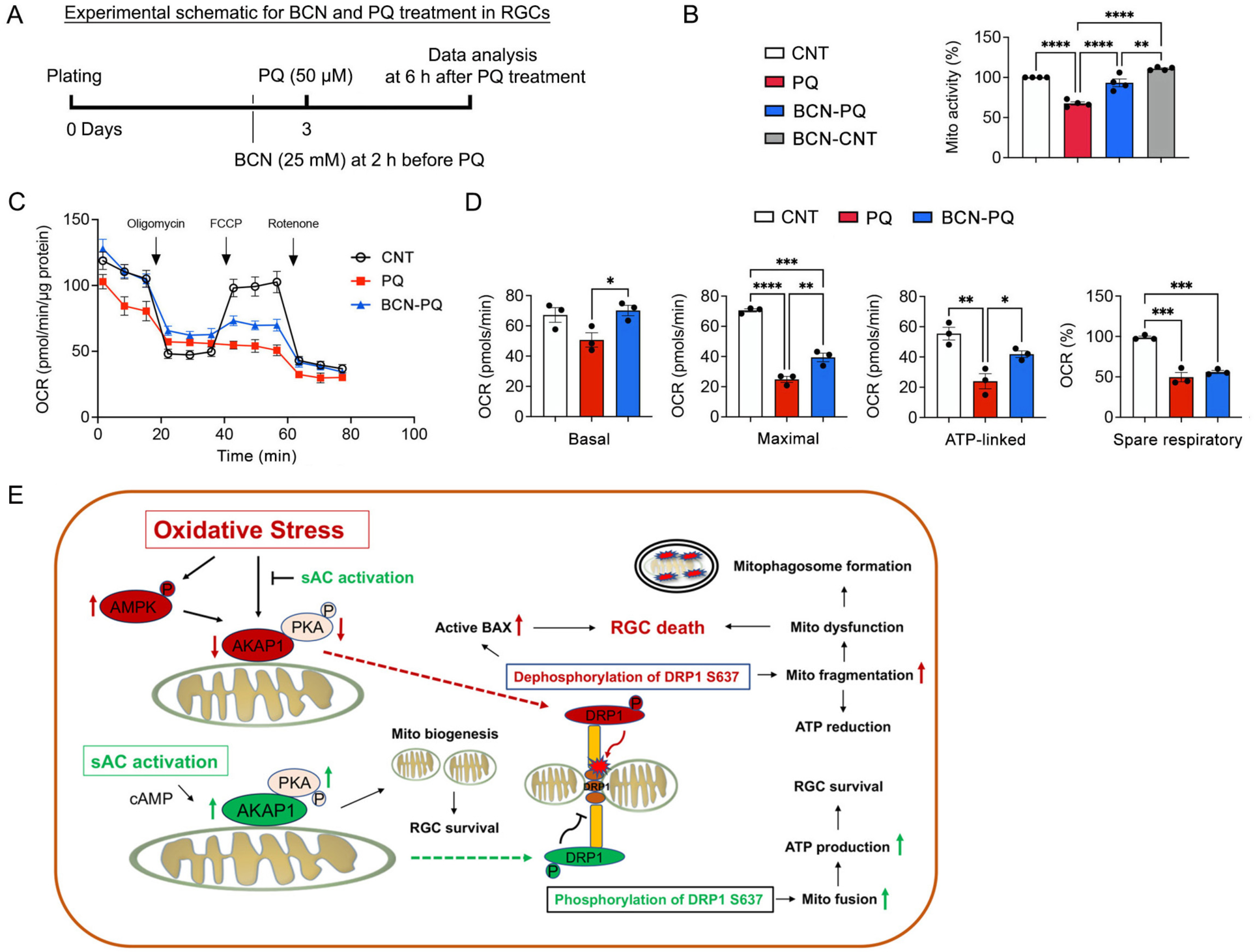
sAC activation promotes mitochondrial bioenergetic function in oxidatively stressed RGCs. (*A*) Diagram for BCN (25 mM) and PQ (50 μM) treatment, sample collection and data analysis. Cultured RGCs were pretreated with BCN for 2 h before PQ treatment. PQ treatment lasted 6 h. (*B*) The mitochondrial activity was reduced after PQ treatment but protected by pretreatment with BCN. (*C*) OCR changes in cultured RGCs treated with PQ (50 μM) for 6 h. Oligomycin A and FCCP were sequentially added at the indicated time point. Basal respiration indicates the starting basal OCR and the value which was set to 100%. Maximum respiration represents the ratio between FCCP uncoupled OCR and basal OCR. (*D*) Quantitative analyses of basal, maximal and ATP-linked respiration, as well as spare respiratory capacity. (*E*) Hypothetical model for protective functions of sAC against oxidative stress and mitochondrial dysfunction in glaucomatous neurodegeneration. The activation of sAC is proposed to mitigate mitochondrial dysfunction, protect RGCs, and alleviates visual impairment caused oxidative stress induced by ischemia/reperfusion. The activation of sAC is suggested to protect RGCs under oxidative stress conditions by promoting mitochondrial biogenesis and OXPHOS activity, while also inhibiting apoptosis. This study establishes a crucial connection between the activation of and the preservation of mitochondrial structure and function in RGCs, offering potential insights into protecting against glaucomatous challenges, particularly oxidative stress. *N* = 3 independent experiments. Error bars represent SEM. Statistical significance determined using one-way ANOVA test. \**P* < 0.05; \*\**P* < 0.01; \*\*\**P* < 0.001; \*\*\*\**P* < 0.0001.

## DISCUSSION

Oxidative stress plays a pivotal role as a regulator in the neurodegeneration of the retina associated with various ocular diseases, including glaucoma. However, little is known about the potential links and molecular mechanisms between sAC activation and mitochondrial protection in RGCs under oxidative stress. In the present study, we demonstrate novel protective roles of sAC activation on RGCs and visual function that counter the effects of oxidative stress. The neuroprotective effects of sAC activation were mediated by promoting expression of AKAP1 and PKAα, enhancing phosphorylation of DRP1 and GSK3β, mitochondrial biogenesis and OXPHOS activity, as well as by reducing AMPK activation, phosphorylation of p38 and BAX activation. In parallel, sAC activation significantly preserved mitochondrial respiratory and structure in RGCs under oxidative stress. Here, we propose that the sAC/cAMP/PKA axis could be a therapeutic target for ameliorating mitochondrial dysfunction and RGC death caused by glaucomatous insults such as oxidative stress.

Increasing the intracellular level of cAMP by electrical activity-mediated depolarization or bicarbonate exposure promotes RGC survival as well as its axon and neurite outgrowth^11, 16^. Of note, the cAMP signaling pathway is linked to the enhancement of PGC1-α activity and the regulation of mitochondrial biogenesis^39, 40^. Moreover, intra-mitochondrial cAMP is likely protective against apoptosis^41^. Indeed, current evidence surprisingly indicates that activating sAC promotes RGC survival and improves visual function by enhancing mitochondrial biogenesis and inhibiting apoptotic cell death. Excessive mitochondrial fragmentation has a crucial role in glaucomatous RGC degeneration^2, 42^. Moreover, inhibiting the activity of DRP1 protects RGCs and/or their axons in glaucomatous DBA/2J and ischemic retina^2, 42, 43^. Under *in vivo* oxidative stress conditions, activating sAC restores the expression level of AKAP1 and PKAα protein and enhances phosphorylation of DRP1 S637 in the retina. Considering that both glaucomatous damage and AKAP1 deficiency result in the dephosphorylation of DRP1 S637 in the retina, leading to excessive mitochondrial fragmentation in RGC somas^2, 23^, our findings suggest that activating sAC has the potential to protect RGCs by modulating cAMP/ PKA/AKAP1-mediated mitochondrial dynamics in response to oxidative stress.

In the current study, we found that sAC activation restored GSK3β S9 phosphorylation under oxidative stress conditions. GSK3β S9 is a substrate of the cAMP/PKA pathway, and PKA has a role in the phosphorylation and inactivation of GSK3β S9^44^. Hence, our findings suggest that the sAC-mediated cAMP/PKA pathway has a critical role in protecting retinal cells by promoting the phosphorylation and deactivation of GSK3β S9 caused by oxidative stress. A previous study reported that GSK3β-dependent phosphorylation of DRP1 S693 promoted mitochondrial elongation in response to oxidative stress^27^. Conversely, blocking GSK3β-mediated phosphorylation of DRP1 S40 and S44 prevented neuronal apoptotic cell death triggered by Aβ-induced toxicity^28^. Intriguingly, recent evidence indicated that reducing the GSK3β level promoted RGC survival and increased the number of neurites^45^. Given the role of GSK3β in the cAMP/PKA/DRP1 axis signaling also associated with mitochondrial elongation^29^, it is likely that sAC-mediated cAMP/PKA/AKAP1/DRP1 axis signaling is linked to protecting the retinal cells through the regulation of GSK3β S9 phosphorylation and mitochondrial elongation, functioning as a protective mechanism against oxidative stress. However, our findings revealed that activating sAC did not significantly enhance mitochondrial fusion activity in the retina under an oxidative stress condition. Therefore, our findings collectively suggest that sAC activation modulates DRP1 or GSK3β phosphorylation, thereby protecting retinal neurons, including RGCs modified by oxidative stress.

Accumulating evidence supports the role of cAMP signaling in regulating mitochondrial biogenesis and OXPHOS^10, 12, 14, 25, 26, 46^. Activation of sAC through bicarbonate or Ca^2+^ increases the cAMP level within the mitochondrial matrix, thereby stimulating OXPHOS activity^47, 48^. OXPHOS dysfunction is linked to the pathogenesis of glaucoma^31^, and evidence suggests that OXPHOS complex I is the primary site of mitochondrial superoxide production by PQ-induced oxidative stress^19^. In our current study, activating sAC enhanced the activities of OXPHOS complexes I and IV in the retina under oxidative stress. This finding aligns with a prior study where inhibiting sAC, achieved through the activation of mitochondria-localize type-1 cannabinoid receptors, resulted in decreased mitochondrial cAMP levels, impaired OXPHOS complex I activity, compromised mitochondrial respiration, and lowered cellular ATP content in neuronal cell cultures^49^. Here, our study provides remarkable evidence that oxidative stress induces abnormal mitochondrial fusion activity, leading to damaged cristae and a subsequent reduction of mitochondrial ATP production in RGCs. More importantly, these findings demonstrate that activating sAC preserves RGC mitochondria by promoting mitochondrial biogenesis and ATP production. Notably, these effects provide significant protection to RGC axon and neurites damaged by oxidative stress. Hence, our findings emphasize a critical role of the sAC/cAMP/PKA signaling axis in mitochondrial protection in RGCs. Activation of cAMP signaling is crucial in mitochondrial biogenesis in mammalian cells^40, 50^ and a deficiency of sAC has been linked to impaired OXPHOS activity^12^. Hence, our findings suggest that the sAC/cAMP/PKA-mediated modulation of mitochondrial biogenesis could protect RGCs by enhancing OXPHOS activity in response to oxidative stress.

As a cellular energy sensor, AMPK is a critical guardian and regulator of metabolic activity that controls normal energy homeostasis^51, 52^ and has a role in mitochondrial fission under energy stress^53^. sAC is involved in regulating AMPK activity, maintaining mitochondrial function, cellular redox balance, and energy homeostasis^37^. Intriguingly, glaucomatous insults, such as elevated IOP and oxidative stress, are linked to AMPK activation in RGCs of the mouse retina and optic nerve by enhancing AMPK phosphorylation^3, 36^. Hence, our findings of activating sAC-mediated inhibition of AMPK activation in stressed RGCs suggest that sAC-regulated AMPK activity contributes to mitochondrial protection in RGCs. Mitophagy or autophagy plays a crucial role in mitochondrial qualify control^54^ and enhanced mitophagosome formation could be protective in RGCs damaged by glaucomatous insults^42, 55^. Indeed, oxidative stress is associated with the reduction of mitophagy by decreasing p62 in glaucomatous retinal degeneration^56^. Here, activating sAC significantly reversed potential reduction in mitophagy by restoring both LC3-II and p62 levels in RGCs under oxidative stress conditions. Together, these findings strongly imply that sAC activation protects RGCs by alleviating mitochondrial stress caused by oxidative stress.

In summary, our study establishes an important link between the sAC/cAMP/PKA axis and mitochondrial regulation in RGC protection and vision restoration after glaucomatous insults such as oxidative stress (Fig. 8E). We suggest that the activation of sAC-mediated mitochondrial protection holds potential as a therapeutic approach for treating glaucoma. Further studies are required to elucidate the potential mechanism(s) underlying sAC/cAMP/PKA-mediated mitochondrial protection against oxidative stress-mediated glaucomatous neurodegeneration.

## MATERIALS AND METHODS

### Animals

C57BL/6J mice (The Jackson Laboratory, ME, USA), classified as adult male and female, were accommodated in enclosed cages, fed with a standard rodent diet, and maintained on a12 h light/12 h dark cycle. The assignment of mice to either experimental or control groups was done randomly. Behavioral response and visual function were studied with 4-month-old 15 male and 15 female mice. Research involving animals in ophthalmic vision conducted by the Association for Research in Vision and Ophthalmology follows protocols approved by the Institutional Animal Care and Use Committee at the University of California, San Diego (USA).

### Pharmacological treatment

NaHCO_3_ was obtained from Sigma-Aldrich (St. Louis, MO, USA). For mouse models of oxidative stress and ischemia-reperfusion, we studied two groups of mice as follows: a group of C57BL/6J mice treated with regular drinking water (*n* = 15 mice for oxidative stress and *n* = 20 mice for ischemia-reperfusion) and a group of C57BL/6J mice treated with drinking water containing 150 mM NaHCO_3_ (*n* = 15 mice for oxidative stress and *n* = 20 mice for ischemia-reperfusion).

### Induction of retinal ischemia-reperfusion by acute IOP elevation

C57BL/6J mice were subjected to anesthesia using a mixture of ketamine (100 mg/kg, Ketaset; Fort Dodge Animal Health, Fort Dodge, IA, USA) and xylazine (9 mg/kg, TranquiVed; Vedeco, Inc., St. Joseph, MO, USA) by intraperitoneal (IP) injection. Induction of ischemia-reperfusion by an acute high IOP elevation was performed as previously described^57^. Briefly, a needle (30-gauge) was introduced into the anterior chamber of the right eye, connected via flexible tubing to a saline reservoir. By elevating the reservoir, the IOP was increased to 70-80 mmHg for a duration of 50 min. Contralateral eyes underwent a sham treatment, involving the insertion of a needle into the anterior chamber without saline injection. Following the removable of the cannula, recirculation commenced immediately, and the IOP returned to normal values within 5 min. Subsequently, mice were anesthetized through an IP injection of a ketamine/xylazine cocktail, as previously described, before cervical dislocation was performed at various time points for tissue preparation after reperfusion: 1 day and 4 weeks. The confirmation of retinal ischemia involved the observation of whitening of the iris and the absence of the red reflex in the retina. IOP was measured with a tonometer (icare TONOVET, Vantaa, Finland) during IOP elevation. Non-IOP elevation contralateral control retinas were used as a sham control.

### Induction of retinal oxidative stress

To induce oxidative stress, mice received IP injection of PQ (15 mg/Kg, Sigma-Aldrich) in saline solution three times during a 1-week period as previously described^23^. Measurements for optomotor response and VEP were assessed at 1 week after PQ treatment.

### Cell culture, and NaHCO_3_ and PQ treatment in vitro

RGCs obtained from 5-day postnatal Sprague-Dawley rats were purified through immunopanning method and cultured in serum-free defined growth medium, following previously established procedures^2^. Approximately 2 x 10^5^, 5 x 10^4^, 1.5 x 10^4^ purified cells were seeded on 60 mm dishes, 24-well plate and 96-well coated first with poly-D-lysine (70 kDa, 10 μg/ml; Sigma-Aldrich), respectively. RGCs were cultured in growth medium containing BDNF (50 μg/ml; Sigma-Aldrich), CNTF (10 μg/ml; Sigma-Aldrich), insulin (5 μg/ml; Sigma-Aldrich), and forskolin (10 μg/ml; Sigma-Aldrich)^2^.

### Western blot analyses

Retinas that were collected were homogenized on ice for 1 min using a modified RIPA lysis buffer (Cell Signaling Technology, MA, USA) as previously described^2, 21^. Proteins (10-20 μg) were run on a NuPAGE Bis-Tris gel (Invitrogen, CA, USA) and transferred to polyvinylidene difluoride membranes (GE Healthcare Bio-Science, NJ, USA). To block membrane, 5% non-fat dry milk and PBS/0.1% Tween-20 (PBS-T) were used for 1 h at room temperature. Subsequently, the membrane was exposed to primary antibodies (sTable 1) overnight at 4°C. Afterward, the membranes underwent three times with PBS-T and were then subjected to incubation with horseradish peroxidase-conjugated secondary antibodies (Bio-Rad, CA, USA) for 1 h at room temperature. The images were captured using a digital imaging system ImageQuant LAS 4000 (GE Healthcare Bio-Sciences Corp., NJ, USA).

### Immunohistochemistry and immunocytochemistry

Immunohistochemical or immunocytochemical staining was performed on wax sections or cultured cells^2^. Sections from each group (*n* = 4 retinas/group) or cells were used for immunohistochemical analysis. To minimize non-specific background, tissues or cells underwent a 1 h incubation at room temperature in 1% bovine serum albumin (BSA, Sigma-Aldrich) in PBS before being exposed to primary antibodies for 16 h at 4°C. Following multiple wash steps, the tissues were subjected to a 4 h incubation with the secondary antibodies (sTable 1) at 4°C, followed by PBS washing. Finally, the tissues were counterstained with the nucleic acid stain Hoechst 33342 (1 μg/ml; Invitrogen) in PBS. Images were acquired with Keyence All-in-One Fluorescence microscopy (BZ-X810, Keyence Corp. of America, IL, USA). Each target protein fluorescent integrated intensity was measured using ImageJ software. All imaging parameters remained the same and were corrected with background subtraction.

### Whole-Mount Immunohistochemistry and RGC Counting

Whole-Mount Immunohistochemistry was performed as previously described ^2^. Retinas were soaked in PBS with 30% sucrose for 24 h at 4°C. Following this, the retinas underwent blocking in a blocking solution composed of PBS containing 3% donkey serum, 1% bovine serum albumin, 1% fish gelatin, and 0.1% triton X-100. Subsequently, the retinas were exposed to primary antibodies (sTable 1) for 3 days at 4°C. Following multiple washing steps, the tissues were treated with secondary antibodies (sTable 1) for 24 h. Images were acquired with Keyence All-in-One Fluorescence microscopy (BZ-X810, Keyence Corp.). RGCs labeled with Brn3a were counted in two zones, specifically the middle and peripheral regions of the retina (corresponding to three-sixths and five-sixths of the retinal radius) (sTable 2).

### Virtual optomotor response analysis

Spatial visual function was assessed using a virtual optomotor system (OptoMotry; CerebralMechanics Inc., AB, Canada) as previously described^21^. To evaluate visual acuity, tracking was observed when the mouse ceased body movement, and only head-tracking movement was examined. The spatial frequency threshold, indicative visual acuity, was automatically determined by the OptoMotry software using a step-wise paradigm based on head-tracking movements at 100% contrast. The spatial frequency initiated at 0.042 cyc/deg and increased progressively until head movement was no longer detected.

### VE) analysis

VEP measurements were conducted following established protocols^21^. The recorded signal was amplified, digitally processed using Veris Instrument software (OR, USA), exported, and subsequently analyzed for peak-to-peak responses. For each eye, the peak-to-peak response amplitude of the major component P1-N1 in IOP eyes was compared to that of their contralateral non-IOP controls. All the recordings were performed with the same stimulus intensity and group comparisons were made based on both amplitude and latency.

### MitoTracker Red staining

Cultured RGCs were pretreated with either saline or NaHCO_3_ (25 mM) for 2 h, followed by exposure to PQ (50 μM) for 24 h. Mitochondria in cultured RGCs were labeled by the addition of a red fluorescent mitochondrial dye (100 nM final concentration; MitoTracker Red CMXRos; Invitrogen, Carlsbad, CA, USA) to the cultures and were maintained for 20 min in a CO_2_ incubator. The cultures were subsequently fixed with 4% paraformaldehyde (Sigma-Aldrich) in DPBS for 30 min at 4°C. Images were acquired with Keyence All-in-One Fluorescence microscopy (BZ-X810, Keyence Corp. of America).

### MTT assay

Mitochondrial activity was assessed using 3-[4, 5-dimethylthiazol-2yl]-2, 5-diphenyl tetrazolium bromide (MTT) in accordance with the manufacturer’s recommendations (Cell Proliferation Kit I; Roche Diagnostics, Basel, Switzerland). In brief, cultured RGCs were pretreated with either saline or NaHCO_3_ (25 mM) for 2 h, followed by exposure to PQ (50 μM) for 24 h. Subsequently, 10 μl MTT stock solution was added to each well, including the negative control. RGCs were treated with 100 μl of solubilization solution. Following an overnight incubation at 5% CO_2_ at 37°C, the absorbance (560 and 690 nm) was examined using a microplate reader (SpectraMax; Molecular Devices, San Jose, CA, USA). Each dataset was obtained from multiple replicate wells within each experimental group (*n* = 3).

### Transmission electron microscopy (TEM)

For TEM analyses, cultured RGCs were pretreated with either saline or NaHCO_3_ (25 mM) for 2 h, followed by exposure to PQ (50 μM) for 24 h. To initiate fixation, the growth medium was removed, and a primary fixative composed of 2% paraformaldehyde and 2.5% glutaraldehyde in 0.15 M sodium cacodylate (pH 7.4) was added at 37°C for 2 min. Following TEM tissue preparation as previously described^21^, an 80kV Tecnai Spirit (FEI; Hillsboro, OR, USA) electron microscope was utilized to capture images with a Gatan 2Kx2K CCD camera at 2.9 nm/pixel. Quantitative analysis involved measuring mitochondrial profile lengths and areas were measured with ImageJ (National Institute of Health)^21^. The number of mitochondria was normalized to the total area occupied by RGC somas in each image, determined using ImageJ and the mitochondrial volume density was estimated using stereology^21^.

### 3D EM tomography

EM tomography analyses were conducted on a FEI Titan Titan High Base electron microscope operated at 300kV with nanoscale spatial resolution and the IMOD package was used for alignment, reconstruction and volume segmentation.as previously described^21^. Measurements of both mitochondria and cristae surface areas, as well as volumes, were conducted within the segmented volumes using IMODinfo. Energy calculations were performed using the biophysical model as previously described^21^.

### OCR analysis

The OCR was measured using an XFe96 analyzer (Agilent, Santa Clara, CA, USA). Cells were pretreated with either saline or NaHCO_3_ (25 mM) for 2 h, followed by exposure to PQ (50 μM) for 6 h. After measuring the basal respiration, oligomycin (2 μg/ml, Sigma-Aldrich), an inhibitor of ATP synthesis, carbonyl cyanide 4-(trifluoromethoxy) phenylhydrazone (FCCP; 1 μM, Sigma-Aldrich), the uncoupler, and rotenone (2 μM, Sigma-Aldrich), an inhibitor of mitochondrial complex I, were sequentially added to measure maximal respiration, ATP-linked respiration, and spare respiratory capacity.

### Statistical analysis

For comparison between two groups that have small number of samples related to a fixed control, statistical analysis was conducted utilizing nonparametrical analysis to one-sample *t*-test. For comparison between independent two groups, a two-tailed Student’s *t*-test was performed. For multiple group comparisons, we used either one-way ANOVA or two-way ANOVA, using GraphPad Prism (GraphPad, CA, USA). Statistically significance was defined as a *P* value below 0.05.

## ACKNOWLEDGEMENTS

We would like to acknowledge a kind gift of *AKAP1^−/−^* mice from Dr. Stefan Strack (University of Iowa). This study was supported in part by National Institutes of Health grants EY031697 to W.-K.Ju and G.A.Perkins, EY034116 to W.-K.Ju, and a P30 EY022589 to Vision Core Grant, as well as an unrestricted grant from Research to Prevent Blindness (New York, NY).

---

**Supplemental Figure 1.**
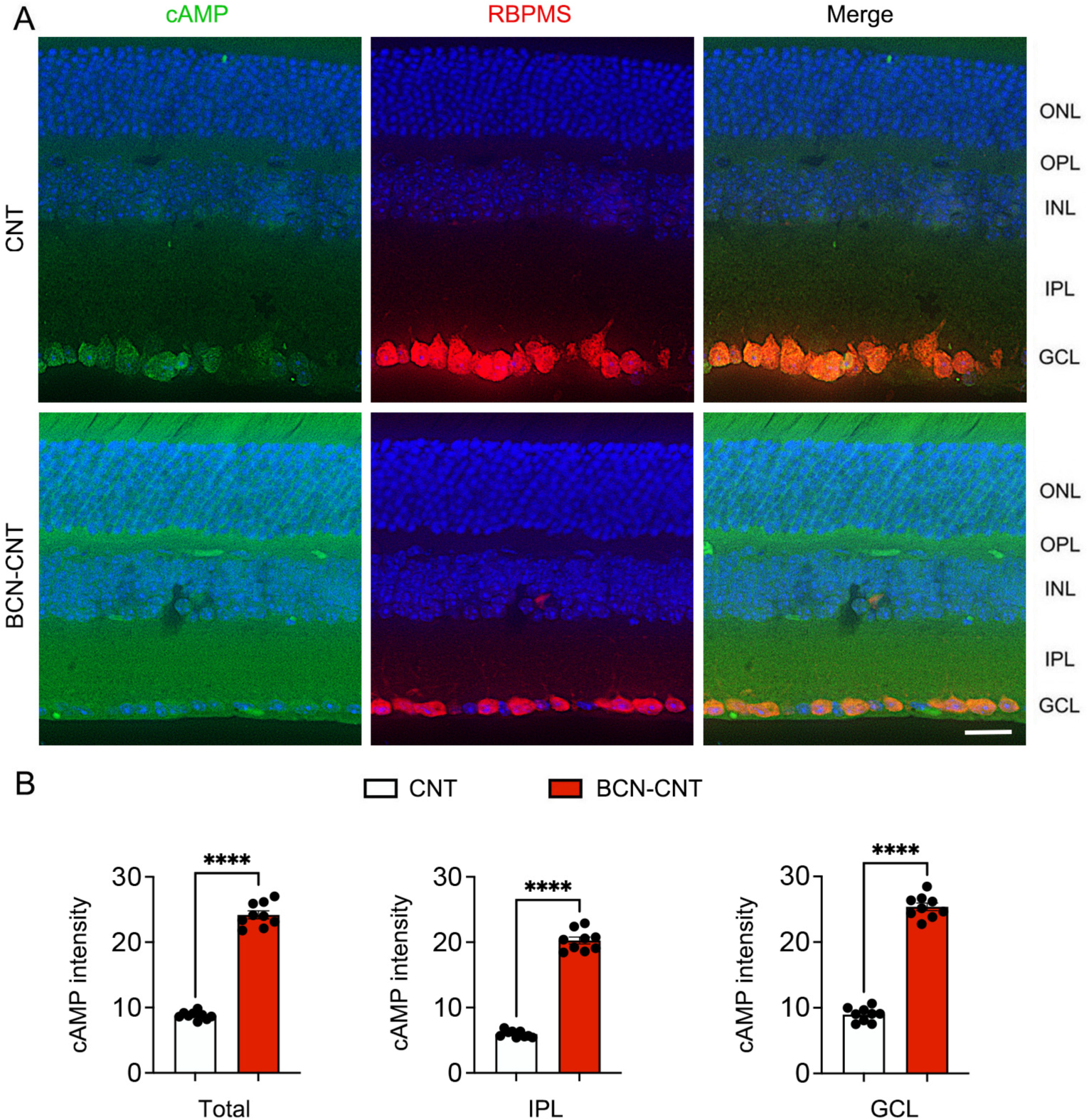
cAMP protein expression in the retina. (A) Representative images showed cAMP (green) and RBPMS (red) immunoreactivities. (B) cAMP immunoreactive intensity in the inner retina. *N* = 3 mice per group. Error bars represent SEM. Statistical significance determined using a two-tailed Student’s *t*-test. \*\*\*\**P* < 0.0001.

**Supplemental Figure 2.**
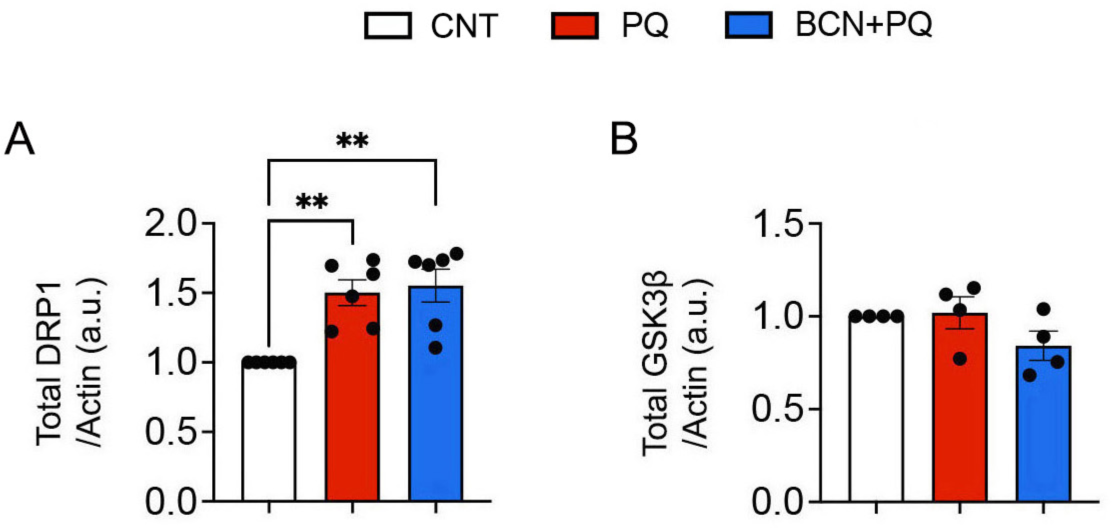
Total DRP1 and GSK3β protein expression in the retina. *N* = 4 to 6 mice per group. Error bars represent SEM. Statistical significance determined using one-way ANOVA test. \*\**P* < 0.01.

**Supplemental Figure 3.**
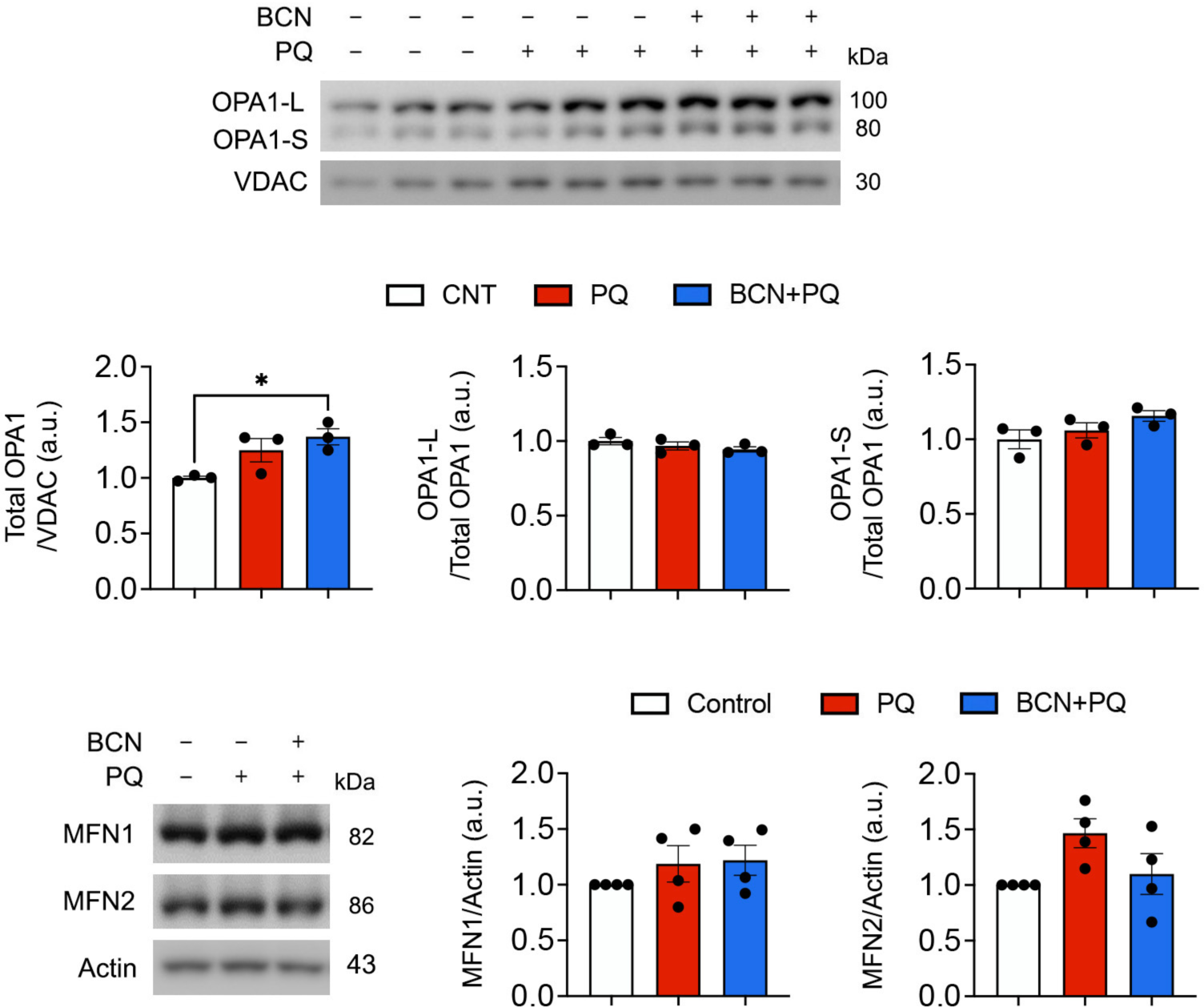
sAC activation did not change the expression level of OPA1 and MFN1 and 2 in the oxidatively stressed retina. (A) OPA1 protein expression in the retina. *N* = 3 mice per group. (B) MFN1 and 2 protein expression in the retina. *N* = 4 mice per group. Error bars represent SEM. Statistical significance determined using one-way ANOVA test. \*\**P* < 0.05.

**Supplemental Figure 4.**
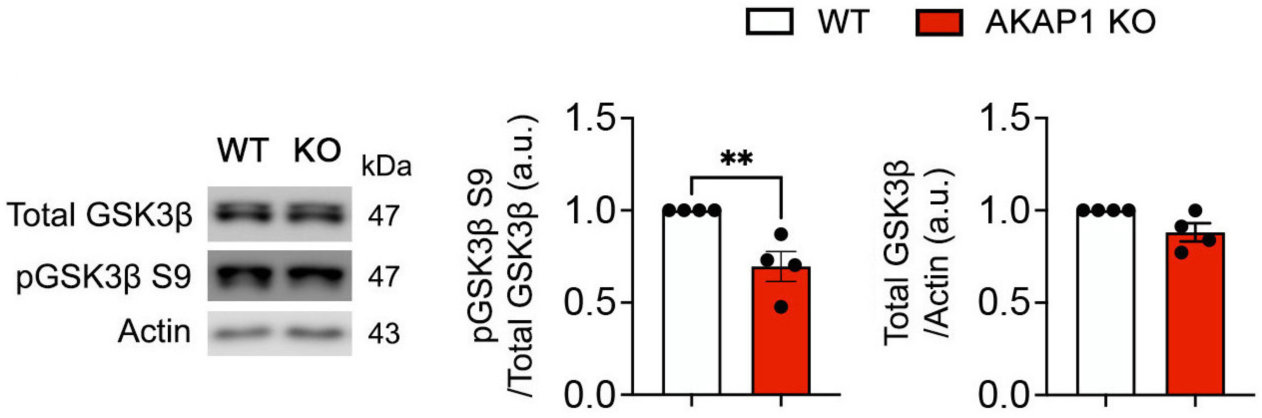
sAC activation induces dephosphorylation of GSK3β S9 in the retina of *AKAP1^−/−^* mice. Total GSK3β S9 and phospho-GSK3β S9 protein expression in the retina. *N* = 4 mice per group. Error bars represent SEM. Statistical significance determined using a two-tailed Student’s *t*-test. \*\**P* < 0.01.

**Supplemental Table 1.** Key Resources.

| REAGENT OR RESOURCES | SOURCE | IDENTIFIER |
| --- | --- | --- |
| Antibodies | Supplier | Catalogue number |
| AKAP1 | Cell Signaling Technology | CST #5203 |
| AMPK | Cell Signaling Technology | CST #5831 |
| Phospho-AMPK | Cell Signaling Technology | CST #2535 |
| β-ACTIN | Millipore | MAB1501 |
| Active BAX | Santa Cruz Biotechnology | sc-23959 |
| BCL-xL | Santa Cruz Biotechnology | sc-8392 |
| Brn3a | Santa Cruz Biotechnology | Sc-8429 |
| cAMP | Abcam | Ab70280 |
| DRP1 | BD Biosciences | #611113 |
| Phospho-DRP1 S616 | Cell Signaling Technology | CST #3455 |
| Phospho-DRP1 S637 | Cell Signaling Technology | CST #4867 |
| GFAP | Invitrogen | 13-0300 |
| GSK3beta | Cell Signaling Technology | CST #12456 |
| Phospho-GSK3b S9 | Cell Signaling Technology | CST #5558 |
| IBA1 | Wako Chemicals | 016-20001 |
| LC3 | Cell Signaling Technology | CST #2775 |
| MFN1 | Abcam | ab57602 |
| MFN2 | Abcam | ab56889 |
| OPA1 | BD Biosciences | #612607 |
| OXPHOS | Invitrogen | #458099 |
| p38 | Cell Signaling Technology | CST #8690 |
| Phospho-p38 | Cell Signaling Technology | CST #4511 |
| p62 | MBL International | PM045 |
| PGC-1alpha | Santa Cruz Biotechnology | sc-13067 |
| PKA | Santa Cruz Biotechnology | sc-28315 |
| IBA1 | Wako | 019-19741 |
| RBPMS | Novus Biologicals | NBP2-20112 |
| TFAM | GeneTex | GTX77852 |
| TUJ1 | BioLegend | 801202 |
| VDAC | Cell Signaling Technology | CST #4866S |
| Goat anti-rabbit HRP | Cell Signaling Technology | 7074 |
| Goat anti-mouse HRP | Cell Signaling Technology | 7076 |
| Alexa Fluor-488 conjugated donkey anti-mouse IgG antibody | Invitrogen | A-21203 |
| Alexa Fluor-568 conjugated donkey anti-mouse IgG antibody | Invitrogen | A-10037 |
| Alexa Fluor-488 conjugated donkey anti-rabbit IgG antibody | Invitrogen | A-21206 |
| Alexa Fluor-568 conjugated donkey anti-rabbit IgG antibody | Invitrogen | A-10042 |
| Chemicals, Reagent and Commercial kits | Supplier | Catalogue number |
| MTT | Roche | 11465007001 |
| MitoTracker Red | Thermo Scientific | M7512 |
| Paraformaldehyde | Sigma-Aldrich | 158127 |
| Paraquat | Sigma-Aldrich | 36541-100mg |
| Sodium bicarbonate | Fisher Scientific | 144-55-8 |
| Seahorse XF Cell Mito Stress Test Kit | Agilent | 103015-100 |
| SuperSignal Chemiluminescent | Thermo Scientific | 34580 |
| Experimental models: Strains | Vendor |  |
| C57BL/6J | Jackson laboratory | Stock No: 000664<br>RRID:IMSR_JAX:000664 |
| AKAP1 KO |  |  |
| Sprague-Dawley Rat | Envigo (Harlan) | N/A |
| Software and Algorithms | Suppliers | Site Link |
| ImageJ | NIH, USA | https://imagej.nih.gov/ij/ |
| Adobe Photoshop | USA | https://photoshop.com |
| Prism 9 | GraphPad Inc. | https://www.graphpad.com/scientific-software/prism/ |

**Supplemental Table 2.** Effect of sAC activation on Brn3a-positive RGC survival in the middle and peripheral retina from mice induced by ischemia-reperfusion.

| Strain | Treatment | Model | Age (Months) | RGC density per retina (RGCs/mm<sup>2</sup>) |  |
| --- | --- | --- | --- | --- | --- |
|  |  |  |  | Middle | Peripheral |
| C57BL/6J | Drinking water | CNT | 4 | 3474 ± 148 | 2619 ± 169 |
| C57BL/6J | Drinking water | EIOP | 4 | 2243 ± 101 | 1669 ± 141 |
| C57BL/6J | NaHCO<sub>3</sub> in drinking water | EIOP | 4 | 3217 ± 60 | 2595 ± 67 |
| C57BL/6J | NaHCO<sub>3</sub> in drinking water | CNT | 4 | 3362 ± 65 | 2530 ± 62 |

All results were reported as means ± SEM. *n* = 5-8 retina wholemounts from 5-8 mice per group. EIOP, elevated intraocular pressure.

## Supplemental Movie Legend

**Supplemental Movie 1.** EM tomography showed that mitochondria were typically elongated with slightly condensed matrix (expanded cristae) in control RGC.

**Supplemental Movie 2.** Oxidative stress produced longer mitochondria, yet fewer in number, and abnormal mitochondrial membranes in PQ-treated RGC.

**Supplemental Movie 3.** sAC activation increased the crista density in PQ-treated RGC.

